# The novel P_II_-interacting protein PirA regulates flux into the cyanobacterial ornithine-ammonia cycle

**DOI:** 10.1101/2020.11.23.395327

**Authors:** Paul Bolay, M. Isabel Muro-Pastor, Rokhsareh Rozbeh, Stefan Timm, Martin Hagemann, Francisco J. Florencio, Karl Forchhammer, Stephan Klähn

## Abstract

Among prokaryotes, cyanobacteria have an exclusive position due to the fact that they perform oxygenic photosynthesis. Cyanobacteria substantially differ from other bacteria in further aspects, e.g. they evolved a plethora of unique regulatory mechanisms to control primary metabolism. This is exemplified by the regulation of glutamine synthetase (GS) via small proteins termed inactivating factors (IFs). Here we reveal another small, 51 amino acid protein, which is encoded by the *ssr0692* gene, to regulate flux into the ornithine-ammonia cycle (OAC), the key hub of cyanobacterial nitrogen stockpiling and remobilization. This regulation is achieved by the interaction with the central carbon/nitrogen control protein P_II_, which commonly controls the entry into the OAC by activating the key enzyme of arginine synthesis, N-acetyl-L-glutamate kinase (NAGK). We suggest that Ssr0692 competes with NAGK for P_II_ binding and thereby prevents NAGK activation, which in turn lowers arginine synthesis. Accordingly, we termed it **<u>P</u>**_II_-**<u>i</u>**nteracting regulator of **<u>a</u>**rginine synthesis (**PirA**). Similar to the GS IFs, PirA accumulates in response to ammonium upshift due to relief from repression by the global nitrogen-control transcription factor NtcA. Consistently, deletion of PirA affects the cell to balance metabolite pools of the OAC in response to ammonium shocks. Moreover, its interaction with P_II_ requires ADP and is prevented by P_II_ mutations affecting the T-loop conformation, the major protein-interaction surface of this signal processing protein. Thus, we propose that PirA is an integrator determining flux into N storage compounds not only depending on the N availability but also the energy state of the cell.

**Importance:** Cyanobacteria contribute a significant portion to the annual oxygen yield and play important roles in biogeochemical cycles, e.g. as major primary producers. Due to their photosynthetic lifestyle cyanobacteria also arouse interest as hosts for the sustainable production of fuel components and high-value chemicals. However, their broad application as microbial cell factories is hampered by limited knowledge about the regulation of metabolic fluxes in these organisms. Our research identified a novel regulatory protein that controls nitrogen flux, in particular arginine synthesis in the cyanobacterial model strain *Synechocystis* sp. PCC 6803. Beside its role as proteinogenic amino acid, arginine is a precursor for the cyanobacterial storage compound cyanophycin, which is of potential interest to biotechnology. The obtained results will therefore not only enhance our understanding of flux control in these organisms, it will also help to provide a scientific fundament for targeted metabolic engineering and hence the design of photosynthesis-driven biotechnological applications.

## Introduction

Nitrogen (N) is one of the key elements of life and needs to be incorporated into biomolecules via assimilatory pathways. Despite an ever-present resource, only a few bacteria can fix dinitrogen (N_2_) and the majority of bacteria relies on the uptake and assimilation of combined N sources from their environment (1–3). To respond to fluctuations in the availability of those N sources, bacteria possess complex regulatory networks to control N uptake as well as the activity of assimilatory enzymes (for reviews see (4–7)). As a prime example, glutamine synthetase (GS), a key enzyme of bacterial ammonium assimilation is tightly regulated in a variety of ways. In *E. coli* and other proteobacteria expression of the GS encoding *glnA* gene is controlled at the transcriptional level by the widespread NtrC/NtrB two-component system (5). Moreover, GS is controlled at the activity level via cumulative feedback inhibition from numerous metabolites related to N and energy metabolism as well as by covalent modification, i.e. adenylylation of the GS subunits. This modification system is operated by a bicyclic modification cascade involving the ubiquitous P_II_ signal transducer protein as regulatory element (reviewed in 8). However, striking differences in comparison to widely accepted paradigms of N assimilation have been revealed in other bacteria, e.g. cyanobacteria.

Cyanobacteria are the only prokaryotes performing oxygenic photosynthesis and play a major role in global biogeochemical cycles (9–11). Nowadays, they receive growing interest as biocatalysts in photo-biotechnological applications, e.g. for the sustainable production of value chemicals and fuels (12–16). To rationally engineer cyanobacteria, i.e. channeling metabolic fluxes to obtain the maximum yield of a desired chemical product, it is of paramount importance to fully comprehend underlying regulatory processes targeting primary metabolism. Albeit our overall understanding of cyanobacterial systems is still fragmentary compared to other well established bacterial models, a few systems have been extensively investigated and include distinctive features. For instance, GS activity is controlled via the interaction with small, inhibitory proteins unique to cyanobacteria (17, 18). These GS inactivating factors (IFs) exclusively control GS activity linearly with their abundance. Moreover, with the global nitrogen control protein NtcA, cyanobacteria use another type of transcription factor to control the expression of genes in response to N fluctuation (19). NtcA belongs to the CRP transcriptional regulator family and commonly works as activator of N assimilatory genes (20–23). During N-limitation, increasing levels of 2-oxoglutarate and the co-activator protein PipX stimulate DNA binding of NtcA (24–26). The interaction between NtcA and PipX is antagonized by the P_II_ protein, which acts as a global multitasking sensor and regulator, adjusting the carbon-nitrogen homeostasis through versatile protein-protein interactions (27, 28). This, for instance, includes the key enzyme for arginine synthesis, N-acetyl glutamate kinase (NAGK), which is activated by complex formation with P_II_ (29).

In addition to the activation of N assimilatory genes, NtcA can also act as a repressor of genes under N limitation. The physiological consequences of simultaneous positive and negative transcriptional regulation are again exemplified by the well-investigated GS regulatory system. During N-limiting conditions, NtcA activates transcription of the *glnA* gene thereby increasing GS abundance and the rate of ammonium assimilation. Simultaneously, enhanced DNA binding of NtcA represses transcription of the genes *gifA* and *gifB* encoding the two known IFs, IF7 and IF17 (30). Thereby, GS activity is tuned in a tradeoff between cellular N demands and relief from the metabolic burden imposed by the glutamate and ATP-consuming GS catalyzed reaction (for a review see 30). Besides *gifA* and *gifB*, only a few other genes appear to be negatively regulated by NtcA. In an attempt to define the entire regulon of NtcA in the unicellular model strain *Synechocystis* sp. PCC 6803 (hereafter *Synechocystis*), Giner-Lamia et al. identified the gene *ssr0692* as another NtcA-repressed candidate (20). It encodes a small protein consisting of 51 amino acids with a high portion of N-rich, positively charged residues, which were shown to be indispensable for protein-protein interaction in case of the GS IFs (18). These distinguishing traits point towards a vital function related to N control similar to the known GS IFs, e.g. as a regulator of a metabolic pathway.

Here we report on the functional analysis of the small protein Ssr0692 in *Synechocystis*. It accumulates in response to ammonium supply and fulfills crucial regulatory roles in cyanobacterial metabolism via the interaction with the P_II_ signaling protein. We suggest that it interferes with the P_II_-dependent activation of NAGK. Consistently, under fluctuating N regimes *ssr0692* mutant strains are impaired in balancing synthesis of arginine and other amino acids associated with the cyanobacterial ornithine ammonia cycle identified recently (32). We therefore named Ssr0692 as **<u>P</u>**_II_-**<u>i</u>**nteracting regulator of **<u>a</u>**rginine synthesis (**PirA**).

## Results

Homologs of the *pirA* gene of *Synechocystis* are frequently present in cyanobacterial genomes and show a high degree of sequence conservation at the amino acid level (**Fig. 1A,B**). With only a few exceptions sequences similar to PirA are absent from genomes of other bacterial phyla (as of July 2020 exceptions are: *Candidatus Gracilibacteria bacterium, Chloroflexaceae bacterium, Flavobacterium sp*. CLA17, *Methylacidiphilales bacterium*). At first glance, this observation suggests a function associated with oxygenic photosynthesis. However, *pirA* has previously been identified as part of the NtcA regulon in *Synechocystis* (20) consistent with two putative NtcA binding motifs located upstream of the transcriptional start site (TSS) (**Fig. 1C**). In promoters that are activated by NtcA, the respective binding motifs are centered close to position −41.5 with regard to the TSS bringing NtcA into a favorable position to promote the binding of RNA polymerase (33). However, the distal motif upstream of *pirA* centers at −48.5, which contradicts the typical position in NtcA-activated promoters. Moreover, the proximal motif centers at −33.5 bp upstream of the TSS, i.e. the first nucleotide is located at position −40 (**Fig. 1C**). This resembles the situation upstream of the *gifA/B* genes, which are also negatively regulated by NtcA (20). Both genes harbor binding motifs that are situated in close proximity or even directly adjacent to the TSS (30), thereby blocking the access of RNA-polymerase. Accordingly, the motif location upstream of *pirA* indicates a negative regulation by NtcA. This assumption is consistent with its downregulation under N limitation, similar to the *gifA/B* genes and in contrast to NtcA-activated genes such as *glnA* or *nrtA* encoding GS and a nitrate transporter component, respectively (**Fig 1D**).

**Figure 1:**
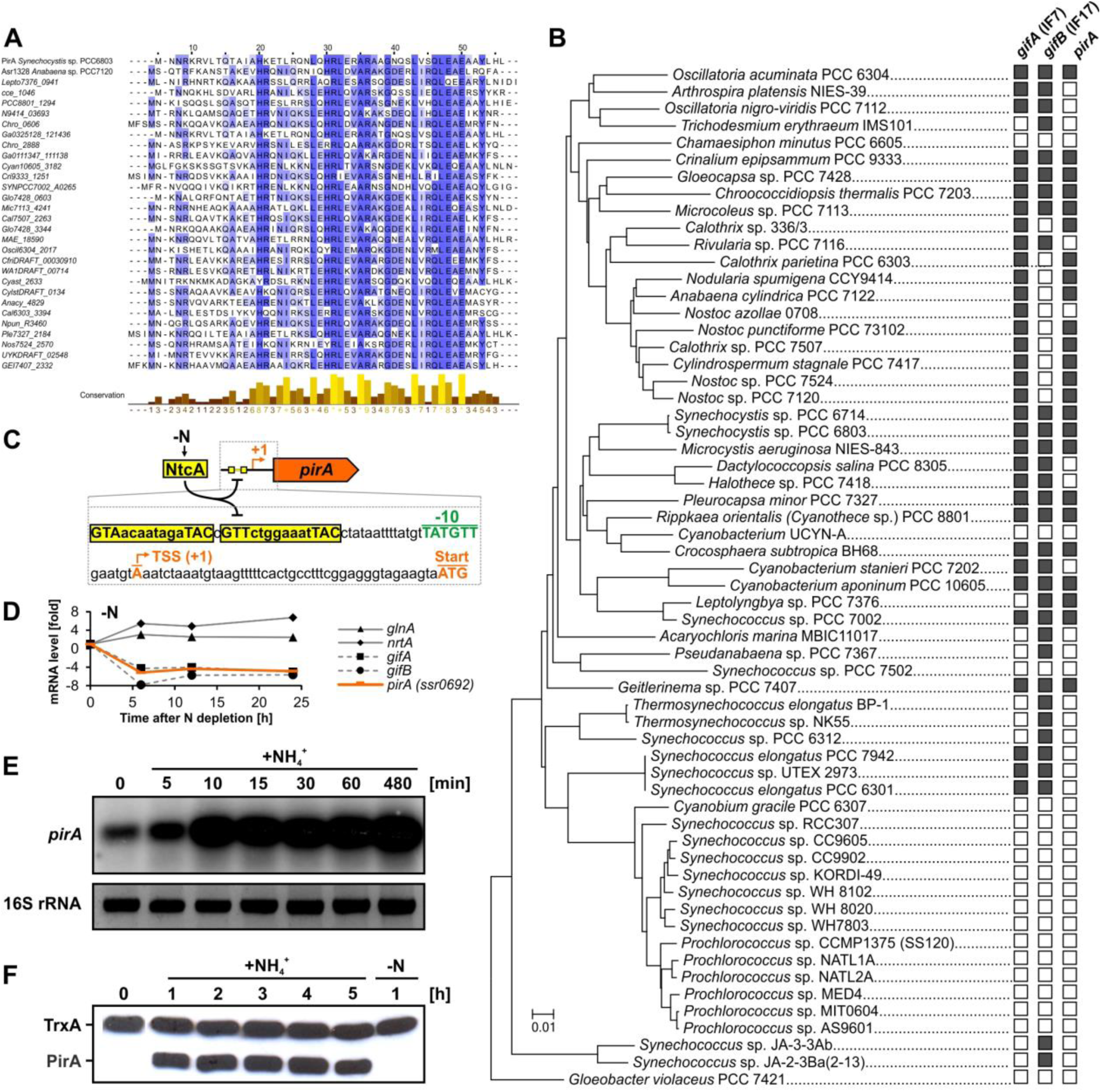
The N-regulated gene *pirA* and its occurrence among cyanobacteria. **A:** Amino acid alignment of randomly selected cyanobacterial PirA homologs. The alignment was made using ClustalW and visualized by using Jalview. **B:** Phylogenetic tree of selected cyanobacteria based on 16S rDNA sequences. The tree was generated with the MEGA7 (82) software package and the Neighbor-Joining method. Please note that we re-used a calculated tree from our previous publication (35) and assigned the presence of genes in the corresponding genomes manually. Gene presence (illustrated by filled rectangle) was investigated using the BLASTP algorithm (83). As reference the amino acid sequences of PirA, IF7 (GifA, Ssl1911) and IF17 (GifB, Sll1515) from *Synechocystis* were used. **C:** Overview of the promoter region upstream of the *pirA* gene in *Synechocystis*. Two putative NtcA binding sites are highlighted. The transcriptional start site (TSS, +1) and the location of the −10 element were extracted from differential RNAseq data (84). **D:** Changes of mRNA levels for several *Synechocystis* genes in response to N limitation. Data were extracted and plotted from previously published microarray data (85). **E:** Northern blot showing transcript accumulation of *pirA* in nitrate-grown *Synechocystis* cells upon addition of 10 mM ammonium chloride. 16S rRNA was used as loading control **F:** Western blot showing changes in PirA protein levels in response to ammonium upshifts and subsequent N depletion. For this a specific, customized antibody against PirA has been raised in rabbit. An antibody against thioredoxin (TrxA) was used to verify equal loading.

### PirA accumulates under N excess and is linked to a function in cyanobacterial N metabolism

Genes that are repressed by NtcA, such as the *gifA/B* genes, show low or even non-detectable transcription under N limitation but are highly expressed in response to excess N supply. To test whether this is also true for *pirA*, we pre-cultivated *Synechocystis* cells in presence of nitrate and analyzed transcript levels after induction of N excess by adding 10 mM ammonium. As expected, the *pirA* mRNA strongly accumulated under these conditions (**Fig. 1E**). To investigate whether this regulatory pattern is conveyed to the protein level, we obtained an antibody specific to the PirA protein. Consistent with the observed transcriptional regulation, the PirA protein also accumulated in response to ammonium upshifts (**Fig. 1F**). Moreover, the protein appeared to have a high turnover due to the fact that it eluded detection shortly after N was depleted. These observations clearly link PirA and its function to cyanobacterial N metabolism.

To investigate the biological function of PirA, knockout and overexpression strains for the *pirA* gene were established in *Synechocystis*. The knockout mutant Δ0692 was generated by replacing the entire *pirA* open reading frame with a kanamycin resistance cassette via homologous recombination. In case of the overexpression strain 0692^+^, a pVZ322 plasmid derivative harboring a transcriptional fusion of *pirA* with the Cu^2+^- inducible *petE* promoter was transferred into *Synechocystis* WT (for a schematic overview of the constructs see **Fig. 2A**). Full segregation of the mutant allele in Δ0692 as well as the presence of the recombinant plasmid in 0692^+^ were verified by PCR (**Fig. 2B**). Subsequent Northern-blots with RNA isolated from cells grown in presence of 1 µM CuSO_4_ confirmed the generated mutants: the overexpression strain showed increased *pirA* mRNA levels compared to WT while in the knockout strain the *pirA* transcript was absent (**Fig. 2C**). Interestingly, even though the mRNA was present and its abundance significantly increased in strain 0692^+^ due to the ectopic expression triggered by Cu^2+^, the PirA protein could not be detected in nitrate-grown cells. However, after adding ammonium, which triggers expression of the native *pirA* gene from the chromosome, increased PirA levels were detectable in strain 0692^+^ compared to the WT (**Fig. 2D**). This, in addition to the confirmed increase at the mRNA level, clearly confirmed that the overexpression construct is operative. Obviously, PirA abundance is not exclusively controlled at the transcriptional level. This observation was further supported by experiments using a *pirA* knockout mutant in which a P*petE*- fused gene copy was introduced. As observed before, the PirA protein could not be detected after adding Cu^2+^ to nitrate-grown cells (**Fig. 2E**). Remarkably, its presence was still N-dependent similar to WT, i.e. it was detectable only after adding ammonium (**Fig. 2E**) even though *pirA* transcription was controlled by P*petE* and hence, exclusively triggered by Cu^2+^. Consequently, these data indicate an additional, posttranscriptional control mechanism, which obviously prevents stable PirA accumulation unless N availability suddenly increases. This again resembles the GS IFs encoded by the *gifA/B* genes, which are tightly regulated at the transcriptional as well as posttranscriptional level (34–36).

**Figure 2:**
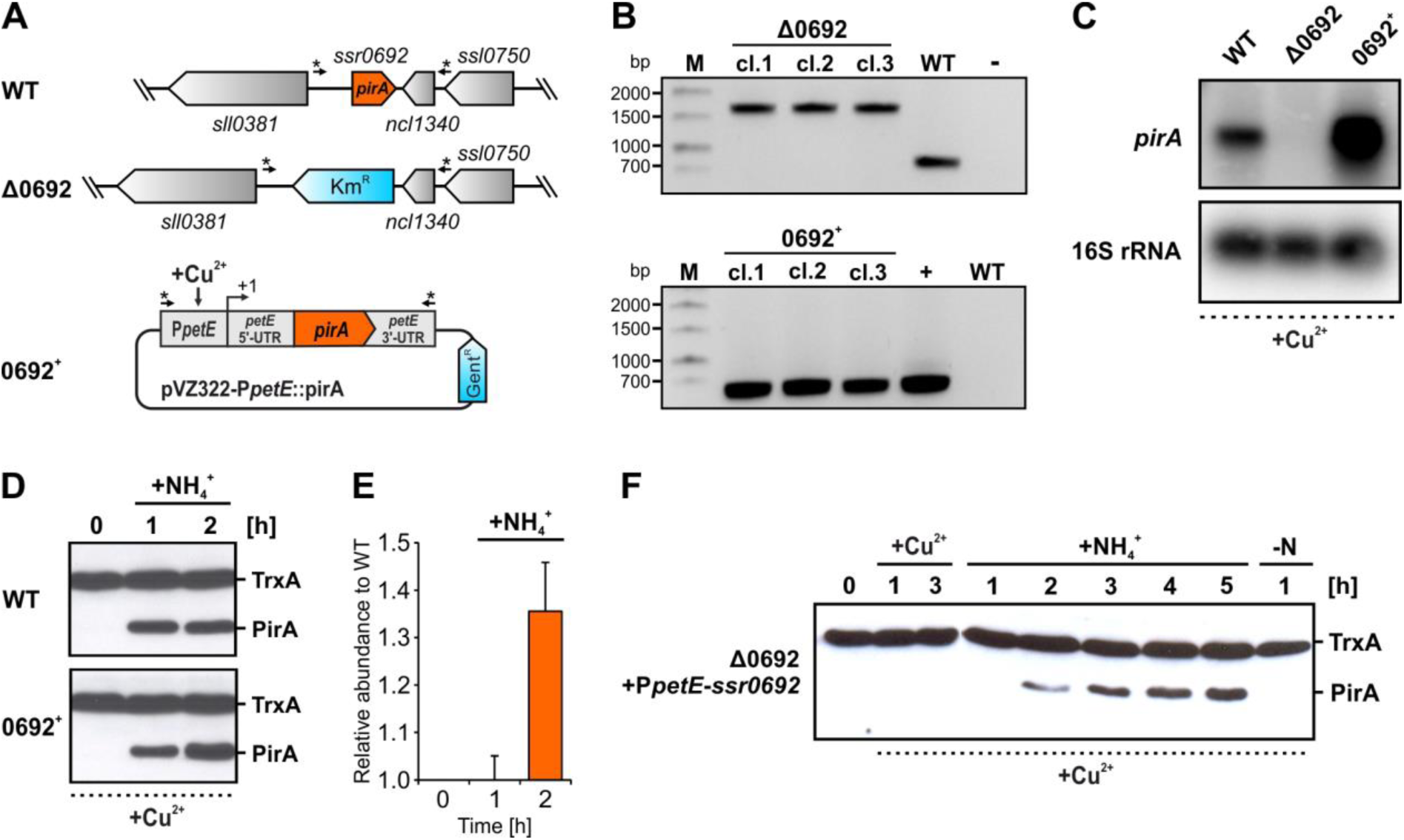
Properties and expression profiles in recombinant strains Δ0692 and 0692^+^. **A:** Schematic view of the *pirA* locus in the WT and in the knockout strain Δ0692 as well as of a pVZ322 plasmid derivative harboring a *pirA* gene copy under control of the Cu^2+^-inducible promoter P*petE* that is present in the overexpression strain 0692^+^. In the Δ0692 knockout strain, *pirA* was replaced by a kanamycin resistance cassette (Km^R^) via homologous recombination. The plasmid enabling ectopic *pirA* expression was introduced into *Synechocystis* WT. The arrows labelled with asterisks indicate the binding sites for primers used to verify the mutants. **B:** PCR verification of the genotype of independently obtained mutant strains. In each case three clones were tested using primer combinations Ssr0692_KO-seg_fw/Ssr0692_KO-seg_rev (in case of Δ0692) or PpetE_fw(XhoI) and Toop_rev(AseI) (in case of 0692^+^). M, marker; bp, base pairs; cl., clone; –, negative control (water as template); +, positive control (purified plasmid as template). **C:** Relative abundance of the *pirA* mRNA, measured via northern blot using sequence specific ^32^P-labelled ssRNA probes. In all cases RNA was isolated from cells grown in presence of 1µM CuSO_4_. **D:** Western blot showing PirA protein levels in cells of the WT and strain 0692^+^, treated with 1 µM Cu^2+^ for 3 hours and afterwards with 10 mM ammonium. Thioredoxin (TrxA) levels verify equal loading. **E:** PirA levels relative to WT. Data were obtained by densitometric evaluation of respective bands using the ImageJ software (86). Data are the mean ± SD of values obtained from two independent western blots, i.e. two biological replicates (independent clones). **F:** PirA accumulation in a Δ0692 strain that was complemented with a *pirA* gene fused to the *petE* promoter. Please note that the data shown here were obtained using a mutant in which the P*petE*-*pirA* construct was integrated into the chromosome, i.e. this strain does not harbor the plasmid derivative shown in panel A.

### PirA plays a critical role upon changes in the C/N balance

Under standard conditions, i.e. with nitrate as sole N source and under ambient CO_2_, at which PirA is not detectable in WT, the *pirA*-manipulated recombinant strains grew similarly to WT as expected (**Fig. 3A,C**). Given the fact that PirA rapidly accumulated in response to an increasing N availability, which suggests a function related to these conditions, it was tempting to speculate whether both recombinant strains show a phenotype, e.g. an affected pigment synthesis/degradation when the N concentration is altered. In order to test this, we cultivated the WT, Δ0692 and 0692^+^ under N oscillating conditions. For that, we inoculated cultures in nitrate-free BG11 and cultivated for three days, which was accompanied by pigment degradation (**Fig. 3B,C**), causing nitrogen-starvation induced chlorosis (37). Cultures of both recombinant strains showed the same behavior as the WT and did not show a non-bleaching phenotype as for instance known for *nbl* mutants that are affected in phycobilisome degradation (38). Consistently, the phycocyanin content was strongly reduced in all cells, measured by the diminished absorption at 630 nm (**Fig 3D**, Day 3). The similar bleaching kinetics of all strains is consistent with the fact that PirA is not detectable under N limitation. Afterwards, the fully chlorotic cells were exposed to consecutive pulses of limited amounts of ammonium (1 mM) to simulate conditions at which PirA is rapidly accumulating and likely important. The re-greening process was monitored by measuring growth as well as whole cell absorption spectra at wavelengths in the range between 400 and 750 nm. While growth recovery was rather similar in all three strains, a clearly altered pigmentation was observed in strain 0692^+^ after these ammonium pulses (**Fig. 3C, D**). Consistent with the visible difference, a lower absorption at 630 nm was detected resulting from a reduced phycocyanin content. These data indicate that the cells are impeded in coping with fluctuating N concentrations and struggle to recover from chlorosis when PirA accumulation is not correctly balanced. This supports the assumption that this small protein plays a crucial role and might participate in regulatory processes that control N metabolism.

**Figure 3:**
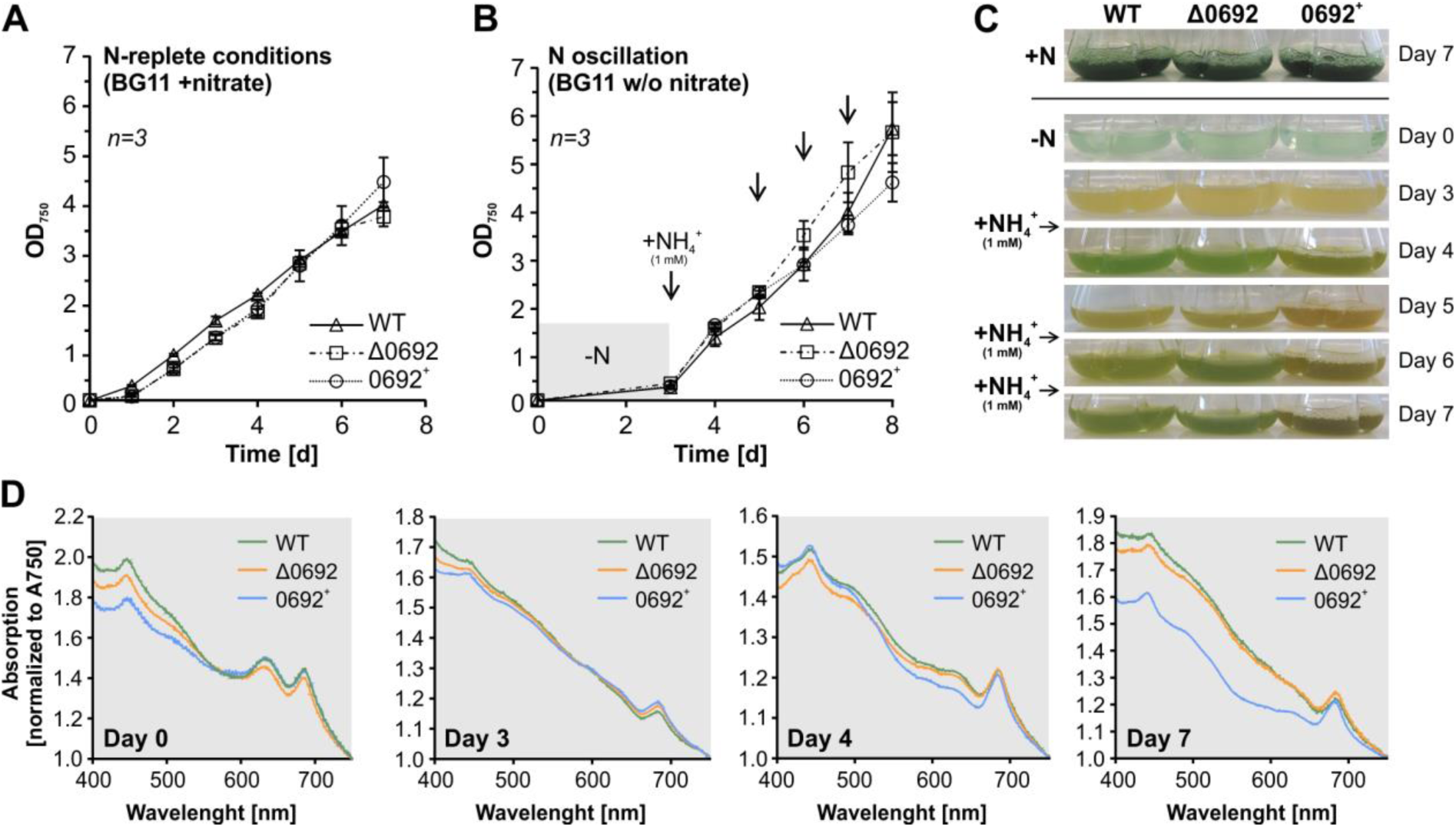
Growth and pigmentation of the WT and the mutant strains Δ0692 and 0692^+^ when N is oscillating. **A, B:** Growth under standard conditions and when ammonium is consecutively added to N starved cultures. Arrows indicate time points at which 1 mM NH_4_Cl was added. Data are the mean ± SD of three independent cultures (including three independent clones of each mutant). **C:** Representative photographs of cultures used in the experiment. Ammonium was added after day 3 and repeated after days 5 and 6. **D:** Whole cell absorption spectra. Values were normalized to A_750_ values.

### Altered PirA abundance affects metabolites of N metabolism

To further examine a potentially regulatory function of PirA, time-resolved quantification of selected metabolites was performed for nitrate-grown cells of the WT and both mutants after addition of 10 mM ammonium. Interestingly, perturbation of PirA levels had a distinctive impact on the accumulation of several key metabolites in *Synechocystis*. Most intriguingly, kinetics of metabolites that are part of or are associated with the recently discovered ornithine-ammonia cycle (32) were strongly affected in strains Δ0692 and 0692^+^ compared to the WT (**Fig. 4**). In general, N upshift triggered a transient accumulation of citrulline, ornithine as well as arginine and aspartate in WT cells similar to previous reports (32). Interestingly, absence of PirA intensified and prolonged the accumulation of these metabolites while overexpression of *pirA* prevented or delayed their accumulation to a significant extend (**Fig. 4**). Moreover, kinetics of glutamine and glutamate, both key amino acids in N metabolism, showed striking differences between the tested strains. For instance, glutamate, which represents the main N-donor in a plethora of pathways was significantly decreased in strain 0692^+^ throughout the experiment. The data clearly indicate that PirA plays a pivotal role in balancing fluxes through or into key amino acids such as arginine. In cyanobacteria, the rate-limiting step of arginine synthesis is controlled by a well-investigated regulatory mechanism, through complex formation of the key enzyme NAGK with the P_II_ protein (28, 29, 39). Moreover, a PII variant with highly increased affinity towards NAGK (PII-I86N) causes constitutive NAGK activation and hence arginine accumulation (40), which was also observed in our study in cells of Δ0692. It was thus tempting to speculate if PirA might interfere at this regulatory node.

**Figure 4:**
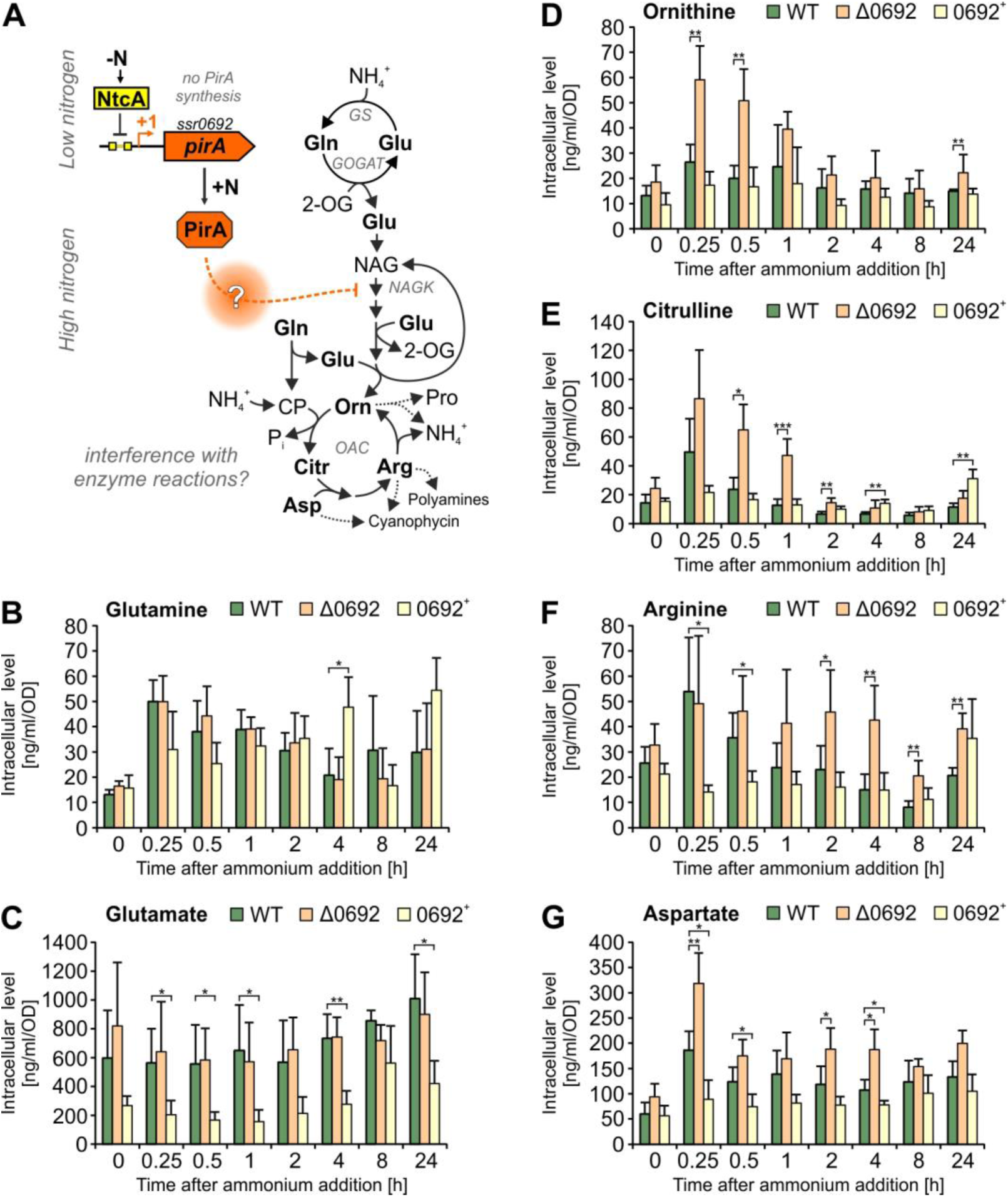
Kinetics of metabolites linked to the OAC cycle in response to ammonium addition. **A:** Simplified overview of metabolic pathways associated with ammonium assimilation and a possible regulatory impact of PirA on certain enzymatic reactions. 2-OG - 2-oxoglutarate, CP - carbamoyl phosphate, GS - glutamine synthetase, GOGAT - glutamine oxoglutarate aminotransferase, NAG - N-acetyl-glutamate, NAGK - N-acetyl glutamate kinase, OAC - ornithine-ammonia cycle. **B-G:** Kinetics of selected metabolites after adding 10 mM ammonium to nitrate-grown cells in the exponential phase. Metabolites were determined by UHPLC-MS/MS after ethanol extraction from cells of the WT, Δ0692 and 0692^+^. Data are the mean ± SD of two independent experiments, each conducted with three biological replicates (independent clones). Significant differences between the strains are labeled and were revealed by one-way analysis of variance (ANOVA; * p<0.05, ** p<0.01, *** p<0.001).

### PirA interacts with the signaling protein P_II_ in an ADP-dependent manner

Recently, PirA was found enriched in pull down-experiments of the signaling protein P_II_ (41). This indicated that PirA may directly interact with the P_II_ protein and thereby exercise a regulatory function similar to other small P_II_ interacting proteins such as PipX (24) or CfrA/PirC that has recently been discovered by two independent laboratories (42, 43). To verify the interaction between PirA and P_II_ from *Synechocystis, in vitro* binding experiments were performed using Bio-layer interferometry (BLI). To this end, recombinant protein variants were expressed in and purified from *E. coli*. His_6_-tagged P_II_ protein was immobilized on a Ni-NTA coated sensor tip and a GST-tagged PirA variant was used as analyte in presence or absence of various effector molecules (**Fig. 5A**). Indeed, complex formation was detected in presence of ADP in a clear concentration dependent manner (**Fig. 5B**). In contrast, no interaction was observed in presence of ATP, mixtures of ATP and 2-OG or when no effector molecule was present. These data unambiguously revealed ADP-dependent interaction between P_II_ and GST-tagged PirA. To test the specificity of the interaction, we performed similar measurements using PirA variants, where the GST-tag was removed by proteolytic cleavage. The small PirA peptide yielded a binding signal that was about six-fold lower than the signal observed for the GST fusion protein (**Fig. 5C**). This agrees well with the expected signal, since the BLI-response depends on the mass changes at the sensor tip (GST-PirA vs. PirA: 31.8 kDa/5.8 kDa = 5.5). Furthermore, providing only the GST-tag (26 kDa) did not result in any detectable signal (**Fig. 5C**), which clearly confirms that P_II_ specifically interacts with PirA in these binding experiments. Since the GST-fusion protein is more accurately to handle and the signal is superior to the isolated PirA peptide, further experiments were performed with GST-tagged PirA.

**Figure 5:**
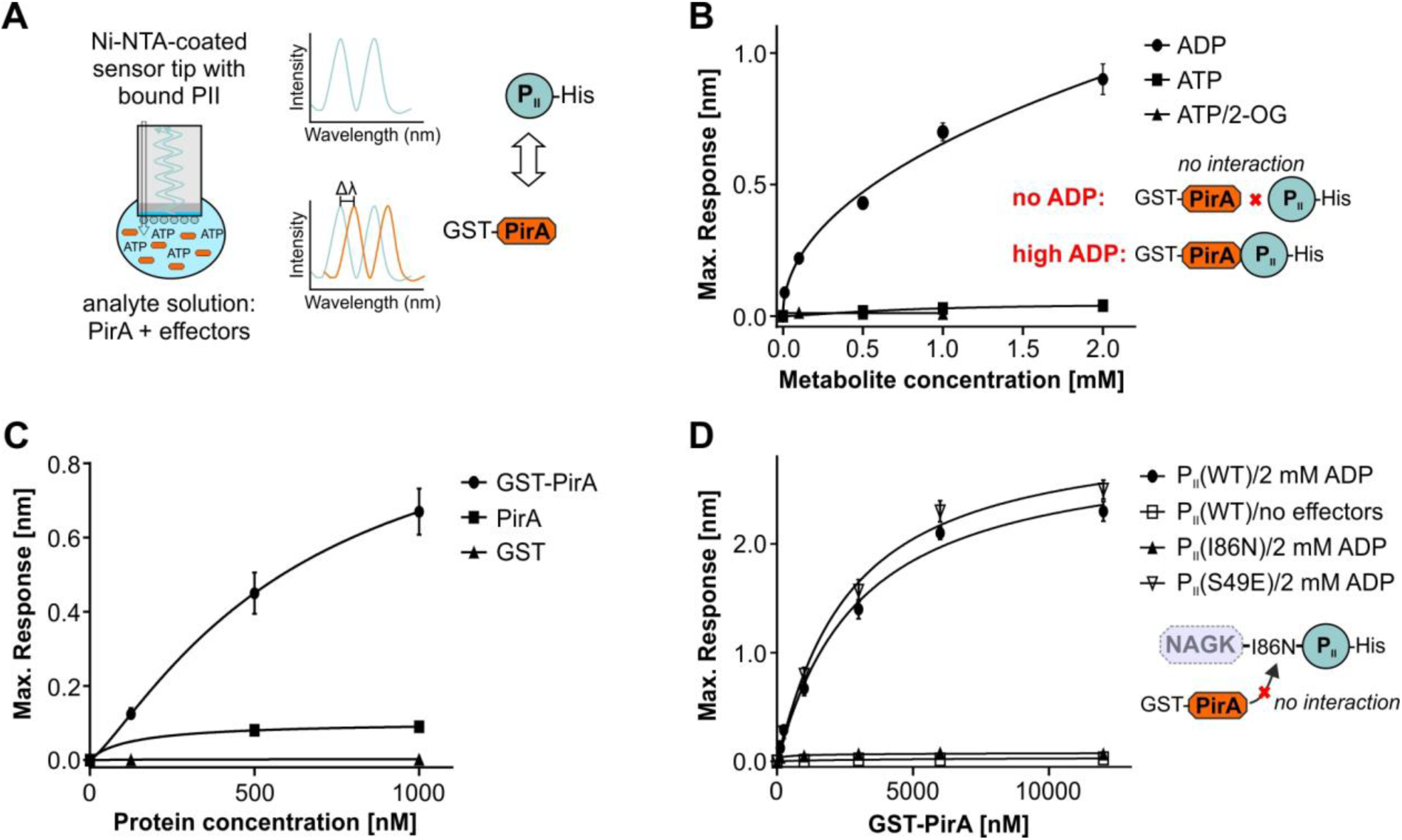
Determination of complex formation between PirA the P_II_ protein measured by Bio-layer interferometry (BLI). **A:** Schematic view of the measuring principle. **B:** Representation of the maximum binding response of P_II_(WT)-His and GST-PirA interaction in the presence of different concentrations of ADP, ATP or ATP/2-OG. **C:** The maximum binding response at different protein concentrations of GST-PirA, tag-free PirA or free GST in the presence of 2 mM ADP. As the binding response is a function of the mass of bound interactor, the response with GST-tagged PirA is correspondingly higher than with isolated PirA peptide. **D:** Representation of the maximum binding response at increasing concentrations of GST-PirA in the absence of effector molecules or in the presence of 2 mM ADP with three different T-loop variants of P_II_. Data are the mean ± SD of triplicate measurements.

To further study the P_II_-PirA interaction, different P_II_ variants were examined. In most cases, interaction of proteins with P_II_ involves the highly flexible T_10_loop structure that can adopt a multitude of conformations (27, 44). Accordingly, a P_II_ variant lacking the T-loop (P_II_(ΔT)-His_8_) was also tested. As expected, no response was observed confirming interaction with PirA via the T-loop (not shown). Moreover, we tested the variant P_II_(I86N), where a single amino acid replacement, Ile86 to Asp86, locks the T-loop in a conformation that promotes constitutive NAGK binding (45, 46). Strikingly, this variant was not able to bind PirA even in presence of 2 mM ADP that otherwise promotes binding to the native P_II_ (**Fig. 5D**). In contrast, the phosphomimetic variant P_II_(S49E), which does not interact with NAGK (47), shows unaffected complex formation with PirA (**Fig. 5D**). Affinity of P_II_(S49E) to PirA was even slightly higher than observed for the native variant (K_D_ values: 2.9 ± 0.34 µM P_II_(WT); 2.5 ± 0.27 µM for P_II_(S49E). Obviously, a conformation of the T-loop that mediates a high P_II_ affinity to NAGK prevents its interaction with PirA. Together with the metabolite profiles showing dysregulated arginine synthesis in the Δ*pirA* mutant, the present data implicate interference of PirA with NAGK regulation through interaction with P_II_.

## Discussion

Arginine serves as proteinogenic amino acid and precursor for the synthesis of polyamines and other N storage compounds such as the cyanobacterial cyanophycin. In bacteria arginine is synthesized either by a linear pathway, e.g. present in *Enterobacteriaceae*, or an energetically more favorable cyclic pathway, where N-acetyl-ornithine reacts with glutamate to yield ornithine and N-acetyl-glutamate, the starting metabolite of the pathway. The latter pathway is widely distributed in nature and also present in cyanobacteria (48, 49). Nevertheless, arginine synthesis requires vast amounts of energy and nitrogen (39) and is thus tightly regulated in bacteria. This is mainly achieved by feedback inhibition of the corresponding key enzymes by the end product arginine. In *E. coli*, this addresses N-acetylglutamate synthase (NAGS), which catalyzes the first step of linear arginine synthesis from glutamate (48). By contrast, in those bacteria harboring the cyclic pathway, the second enzyme NAGK is feedback inhibited by arginine (49).

### A novel player in the distinctive regulation of arginine synthesis in cyanobacteria

In cyanobacteria NAGK is target of a molecular regulatory mechanism that involves complex formation with the signal transduction protein P_II_ (29, 47). This interaction diminishes feedback inhibition of NAGK by arginine and, hence, boosts the metabolic flux towards the end product (29). Its importance for the control of cyanobacterial metabolism is supported by the fact that this mechanism is widely present in oxygenic phototrophs such as plants (50, 51) and microalgae (52) but appears to be absent from other bacteria (for an overview see (53). Regarding arginine metabolism the uniqueness of cyanobacteria within the prokaryotes is also exemplified by the recent discovery of active cycling between ornithine and arginine via an ornithine-ammonia cycle (OAC), similar to the known ornithine-urea cycle (OUC) that is present in terrestrial animals but typically absent from bacteria (32).

Here we propose the small cyanobacterial protein PirA as a novel key regulator in the cyanobacterial arginine synthesis pathway and, hence, also the OAC. Our data confirm PirA accumulation under N excess, in particular when ammonium is added. This accumulation is obviously required to adjust a certain pool of metabolites that are part of the OAC, including arginine. Mechanistically, we propose a model where PirA competes with NAGK for the P_II_ protein (**Fig. 6**). In response to ammonium addition, the high levels of PirA accumulation interfere with complex formation between P_II_ and NAGK, thereby mitigating the activation of NAGK and preventing an over-accumulation of arginine. Surprisingly, detectable PirA accumulation only occurred under ammonium shock, even when *pirA* mRNA was transcribed independently from N and energy status of the cell. A similar situation can be encountered with the GS IFs, which were shown to be degraded by metalloproteases when not bound to GS (34). Thus, it is tempting to speculate that PirA accumulation can only occur when bound to PII.

**Figure 6:**
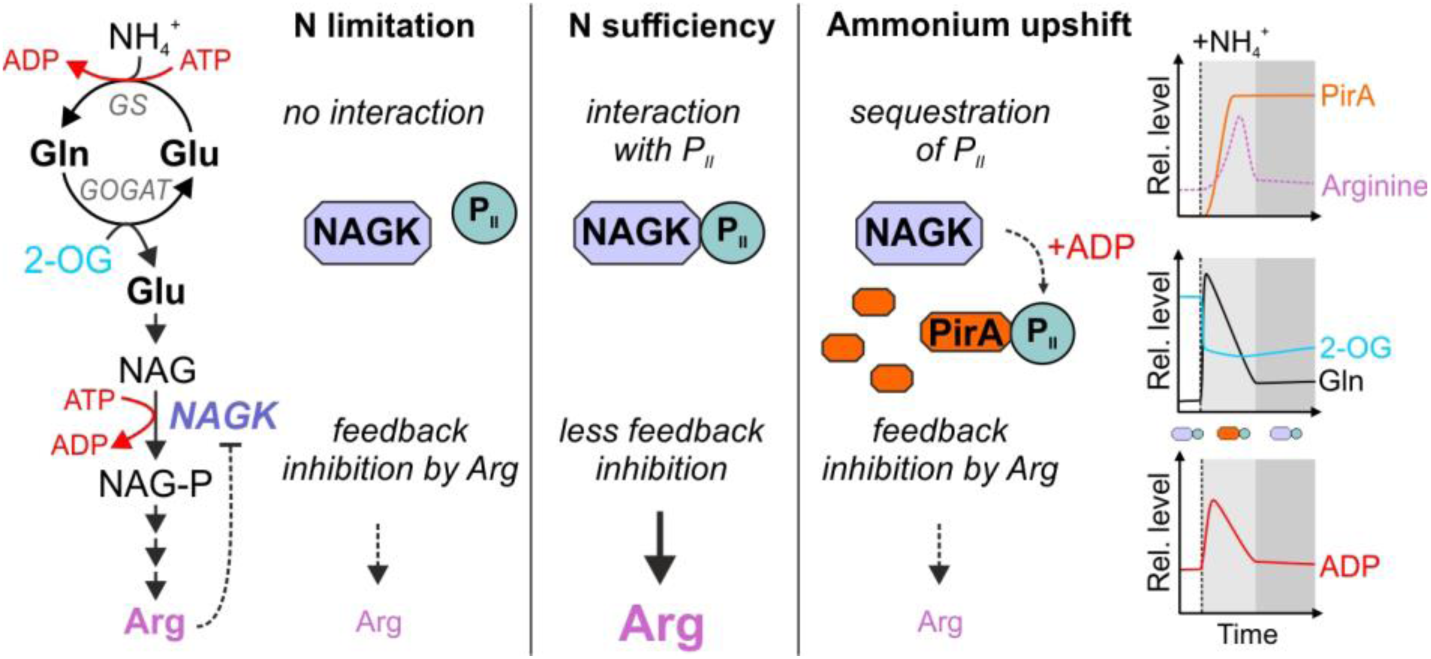
Anticipated model of PirA function. Metabolite kinetics have been approximated based on available literature data (32, 35). Upon shifts in the ammonium concentration PirA accumulates via 2-OG dependent de-repression of the *pirA* gene. The gene product is presumably required to slow down ATP-consuming synthesis of arginine. This could be achieved by ADP-dependent sequestration of P_II_ protein bound to NAGK which is required to diminish feedback inhibition of the enzyme and in turn activate arginine synthesis. The sequestration of P_II_ results in stronger arginine feedback inhibition of NAGK diminishing energy consumption and flux into arginine. After metabolic reorganization (e.g. by inactivating glutamine synthetase activity and decreasing ATP consumption), ADP levels may fall below a critical level preventing interaction between PirA and P_II_. Accordingly, a higher fraction of the P_II_ pool will again interact with and activate NAGK, which in turn results in elevated arginine synthesis.

### PirA integrates N- and energy sensing

Remarkably, PirA only forms a complex with P_II_ in the presence of ADP. Accordingly, the pivotal stimulus for the suggested regulatory mechanism is not only the N status, which is mainly sensed by 2-OG in cyanobacteria similar to other prokaryotes (54, 55). The 2-OG level determines the affinity of NtcA to its binding motif (21, 26), which therefore also determines PirA accumulation. Moreover, PirA was found to transiently but strongly accumulate during low C supply (56). This is consistent with the de-repression of the *pirA* gene by the dissociation of NtcA from its binding motif upstream since low C supply, i.e. a low C/N ratio leads to a decreased 2-OG pool (57). The interaction between PirA and P_II_ strongly responds to ADP as another signal and hence, also depends on the cellular energy status. These observations resemble another small P_II_-interacting protein, PipX, whose interaction with P_II_ is enhanced by ADP as well (58, 59). PipX functions as co-activator of the transcription factor NtcA and is required for the 2-OG-dependent DNA binding and transcriptional activation of genes (24). In presence of high ADP levels, PipX preferably interacts with P_II_, which attenuates NtcA activity and in turn leads to de-repression and upregulation of the *pirA* gene, similar to the *gif* genes encoding the GS IFs (30).

Interestingly, *in vitro* data showed that the NAGK-P_II_ interaction is also ADP-sensitive. While ADP does not entirely prevent complex formation, it increases dissociation of NAGK from ADP-bound P_II_ (45). However, detailed analysis of NAGK-P_II_ complex formation demonstrated that this interaction is mainly tuned by 2-OG and not the energy status of the cell (58, 60). It should be kept in mind that the sensing properties of P_II_ are influenced by the binding partner in such a way, that for certain targets, small fluctuation in the ADP/ATP rations are sensed (e.g. P_II_-PipX complex formation) whereas in other cases, fluctuation in the 2-OG levels are perceived (e.g. P_II_-NAGK interaction). In light of these data, PirA appears to amplify the energy-dependent signal in P_II_-mediated activation of NAGK: by sequestering P_II_ under conditions of low 2-OG and high ADP-levels, PirA may shift the equilibrium of P_II_-NAGK complex towards non-complexed NAGK when the cell experiences energy limitation due to high GS activity in consequence of ammonium addition.

Depending on the detailed structure of the complexes, P_II_ phosphorylation either abrogates interaction with its targets, as demonstrated for NAGK (29), or has no effect on them, as in the case of PII-PipX interaction (33). The phosphomimetic variant P_II_(S49E) showed slightly enhanced interaction with PirA as compared to P_II_(WT). In a similar way, P_II_ interacts with PipX irrespective of Ser49 modification, but nevertheless, the interaction is T-loop dependent (24). The major interaction surface is at the proximal part of the T-loop, whereas the tip region with the critical Ser49 residue does not participate in complex formation. From the Ser49-independent mode of PirA-P_II_ interaction it can be concluded that the PirA-P_II_ interaction could also take place with phosphorylated P_II_ that does not bind to NAGK. In any case, the interaction of P_II_ with NAGK or PirA appears to be mutually exclusive since both PII-interaction partners require the T-loop in different conformation for interaction. The competition between NAGK and PirA for P_II_ is further illustrated by the inability of PirA to bind P_II_(I86N). This variant adopts a constitutive NAGK-bound like structure of the T-loop (45, 46). Accordingly, the lack of interaction with P_II_(I86N) also agrees with the strong *in vivo* activation of NAGK by this variant (40). However, for a more detailed understanding of the mechanism, by which PirA affects P_II_ signaling, functional biochemical studies and structural analyses are required.

### Further implications of PirA beyond arginine synthesis

PirA potentially fulfills several other functions since it appears to feature additional interaction partners, as suggested by previous high-throughput yeast-two hybrid analyses. Among them are NADH dehydrogenase subunit NdhH, DNA polymerase I PolA, the ATP-dependent helicase PcrA as well as several unknown and hypothetical proteins (61). Moreover, *pirA* overexpression was shown to significantly improve butanol tolerance in *Synechocystis* (62). As solvents like butanol compromise cell membrane integrity resulting in loss of proton motive force, the cell increasingly accumulates reactive oxygen species (ROS) through enhanced respiration in an attempt to catch up on lost ATP (63, 64). Given that PirA appears to protect adenylate energy pools, an interaction with players involved in processes that consume large quantities of nucleotides, e.g. DNA replication, appears meaningful. Moreover, overexpression of PirA might result in reduced ROS production as the cell retains sufficient amounts of ATP.

The occurrence of PirA homologs in other cyanobacterial strains does not coincide with a certain lifestyle or phylogeny. For instance, the *pir*A-homolog *asr1328* of the filamentous, diazotrophic cyanobacterium *Anabaena* sp. PCC 7120 was shown to be negatively regulated by NtcA as well (65). Similar to the GS IFs, gene annotations in a few strains such as *Thermosynechococcus elongatus* or *Synechococcus sp*. JA-3-3Ab suggest that alternative versions of PirA with an extended N-terminus might also exist (**Supplementary Fig. S1**). Nevertheless, the existence of those proteins has not been experimentally shown yet, and hence, false annotations cannot be excluded. This assumption is supported by the fact that on the one hand only a few of those elongated sequences could be found in protein databases, which on the other hand only show partial similarity. Therefore, the function of these N-terminal extensions remains elusive to date. In addition, it is worth noting that PirA is, similar to the GS IFs, completely absent in marine picocyanobacteria (**Fig. 1**). In accordance, these clades lack several salient features of N-sensing and utilization which are widespread among cyanobacteria. For instance, P_II_ is not subject to phosphorylation in *Prochlorococcus* (66, 67), and both *Prochlorococcus* and *Synechococcus* genera are incapable of cyanophycin synthesis and lack several OAC cycle genes (68). These genome-streamlined strains occur in oligotrophic realms of the ocean with hardly any fluctuation in nutrient supply (69). Thus, it is compelling to speculate that PirA-mediated short-term adjustment of arginine synthesis to the N and energy status of the cell is not required in such habitats.

## Material & Methods

### Strains and growth conditions

A *Synechocystis* sp. PCC 6803 strain originally obtained from N. Murata (Japan) was used as wild type. Cells were grown in BG11 medium (70) depleted in Cu^2+^ ions and supplemented with 10 mM TES buffered at pH 8.0. The sole N source in that medium is nitrate at a concentration of 17.64 mM (stated as nitrate-grown). Cultivation was performed in buffled Erlenmeyer flasks in presence of ambient CO_2_ under constant illumination (white light, 50µE), at 30°C, 75% humidity and 150 rpm. Each recombinant strain was isolated on BG11-agar plates and maintained in medium containing either kanamycin or gentamycin at a concentration of 50 µg/ml and 2 µg/ml, respectively. Prior to the experiments investigating the impact of altered PirA abundance the cultures were supplemented with 1 µM CuSO_4_.

### Mutant strain generation

To knockout the *ssr0692* (*pirA*) gene, the upstream and downstream region of *pirA* were amplified from *Synechocystis* gDNA using primer combinations Ssr0692upst_fw/Ssr0692upst_rev and Ssr0692downst_fw/Ssr0692downst_rev, respectively (all primers are listed in **Supplementary Table S1**). The kanamycin resistance cassette was amplified from a customized construct obtained by gene synthesis using primers KmR_fw and KmR_rev. The synthesized construct harboured the *aphII* gene and its promoter which were originally obtained from pUC4K (Amersham). In addition, an *oop* terminator was introduced downstream of *aphII*. All amplicons had short fragments of sequence complementarity and were fused via polymerase cycling assembly (PCA) using primers Ssr0692upst_fw/ Ssr0692downst_rev. The resulting construct was introduced into pJET1.2 (Thermo Scientific) and used to transform chemically competent *E. coli* DH5α. After isolation of the pJET_ssr0692_KmR_KO plasmid from *E. coli*, the knockout construct was introduced into *Synechocystis* WT by natural transformation and homologous recombination into the chromosome. To enable overexpression of *pirA*, the 5’ UTR and 3’UTR of the *petE* gene were amplified from *Synechocystis* gDNA using primers PpetE_fw(XhoI)/5’petE_ssr0692_rev and 3’petE_ssr0692_fw/Toop_rev(AseI). The *pirA* coding sequence was amplified from *Synechocystis* gDNA using primers ssr0692_fw/rev. All amplicons were fused via PCR and introduced into the broad-host range plasmid pVZ322 via restriction digestion and ligation into XhoI/AseI sites. The recombinant plasmid pVZ322-PpetE:*ssr0692*, obtained after transformation of and purification from *E. coli* DH5α, was introduced into *Synechocystis* WT via electroporation. All strains and constructs were verified by PCR and Sanger sequencing.

To generate the alternative Δ0692/P_*petE*_-ssr0692 strain (chromosomal integration) a DNA fragment containing ssr0692 gene including the sequence coding for six histidine residues was amplified by PCR using genomic DNA and primers ssr0692.KpnI and ssr0692.HisBamHI. This fragment was cloned into *Kpn*I-*BamH*I digested pPLAT plasmid(34), a pGEM-T derivative containing a 2 kb region of the non-essential *nrsBACD* operon (71). The *petE* promoter was amplified by PCR using genomic DNA and primers P*petE*.KpnI.1 and P*petE*.KpnI.2 and cloned into *Kpn*I site of *ssr0692*-containing pPLAT. Finally, a Kmr CK1 cassette from pRL161 (72) was cloned in the *BamH*I site of pPLAT, generating pPLAT-P*petE*.ssr0692. This plasmid was used to transform the Δ0692 strain. All generated strains and their properties are given in **Supplementary Table S2**.

### N-oscillation experiment

Wild type and mutant strains where inoculated in triplicates at OD_750_ = 0.1 in 3-buffled 100 ml flasks in 20 ml N-depleted BG11 supplemented with 1µM CuSO_4_. For comparison, the same strains were also cultivated in standard BG11 medium containing 17.64 mM nitrate. After 3 days, the N-starved cells where supplemented with 1mM NH_4_Cl, a step which was repeated after 5 and 6 days. Whole cell spectra of cell suspensions were conducted with a Cary 300 UV/Vis Spectrophotometer (Agilent).

### RNA extraction and Northern blots

Cells for northern blot analysis were harvested by rapid filtration on polyether sulfone filters (pore size 0.8 µm, PALL). Filters were immediately resolved in 1 ml PGTX solution (73) and frozen in liquid N_2_. RNA extraction was conducted as previously described (74). For northern blots, 3 µg of RNA were separated on MEN-buffered 1.5% agarose gels that provided denaturing conditions by 6% formaldehyde using an RNA sample loading buffer containing a final concentration of 62.5% (v/v) deionized formamide (Sigma-Aldrich). Afterwards, RNA was transferred via capillary blotting to an Amersham Hybond N^+^ nylon membrane (GE healtcare) and cross-linked with 1250 µJ in a UVP crosslinker (Analytik Jena). To specifically detect the *pirA* transcript, the RNA-mounted nylon membrane was hybridized with a complementary, α-^32^P-labeled ssRNA probe that was generated by *in vitro* transcription using the MAXIscript® T7 Transcription Kit (Thermo Fisher Scientific). As transcription template a DNA fragment obtained by PCR with primers Ssr0692_T7_fw and Ssr0692_rev was used. Subsequently, Fuji BAS-IIIS imaging plates were exposed to the membranes and read out by an Amersham Typhoon laser scanner (GE Healthcare). As loading control, the same membranes were hybridized with ssRNA probes complementary to the 5S rRNA, which were generated in the same way using primers 5sRNA_fw/rev (**Supplementary Table S1**).

### Anti-PirA antibody production

A DNA fragment encompassing the *pirA* ORF was amplified by PCR from *Synechocystis* genomic DNA, using the oligonucleotides Ssr0692ORF-fw and Ssr0692ORF-rv. This fragment was cloned into pET24a(+) plasmid *NdeI*-*Xho*I digested (Novagen) to generate pET24-Ssr0692 plasmid. Exponentially growing *E. coli* BL21 cells transformed with pET24-Ssr0692 were treated with 1 mM of isopropyl ß-D-1-thiogalactopyranoside for 4 h. For purification of PirA-His_6_ protein, cells were collected, resuspended in buffer A (20 mM sodium phosphate, pH 7.5, 150 mM NaCl, 5 mM imidazole) with 1 mM phenylmethylsulfonyl fluoride (PMSF) and disrupted by sonication. The lysate was centrifuged at 18.000 x *g* for 20 min. PirA-His_6_ from the supernatant was purified by Ni-affinity chromatography using HisTrapHP column (GE Healthcare) and following the manufacturer’s instructions. Elution was performed with a linear gradient (5-500 mM imidazole) in buffer A. Fractions with PirA-His_6_ were pooled, concentrated using Centrifugal Filter Units (Amicon Ultra-15 3 kDa) (Millipore), and subjected to gel filtration chromatography using a Hiload 16/60 Superdex 75 gel filtration column (GE Healthcare) running on an AKTA FPLC system. Fractions containing purified PirA-His_6_ protein were pooled, concentrated and quantified in a NanoDrop 1000 spectrophotometer (Thermo scientific) using the extinction coefficient of PirA-His_6_ calculated with ExPASy-ProtParam tool. Anti-PirA antiserum was obtained according to standard immunization protocols by injecting 1.5 mg of purified PirA-His_6_ protein in rabbits.

### Preparation of crude extracts and Western blot analysis

For the analysis of proteins abundance, 2 U OD_750_ were harvested and resuspended in 80 µl of 50 mM HEPES-NaOH buffer (pH 7.0), 50 mM KCl, 1 mM phenylmethylsulfonyl fluoride (PMSF). Crude extracts were prepared using glass beads as previously described (75). For western blot analysis, proteins were fractionated on 15% SDS-PAGE (76) and transferred to nitrocellulose membranes (Bio-Rad). Blots were blocked with 5% (w/v) non-fat dry milk (AppliChem) in PBS-Tween 20. Antisera were used at the following dilutions: Anti-PirA (1:5000) and anti-TrxA (1:10000)(77). The ECL Prime Western Blotting Detection Reagent (GE Healthcare) was used to detect the different antigens with anti-rabbit secondary antibodies (1:25000) (Sigma-Aldrich).

### Metabolite analysis

For metabolite analysis, cells were grown in BG11 containing the standard nitrate amount until reaching OD_750_ ∼0.8. Cells were harvested shortly before and after the addition of 10 mM ammonium chloride by centrifugation of 2 ml culture at 17,000 x *g* for 1 minute. Supernatant was discarded and pellets were snap frozen in liquid N. Metabolite extraction was performed by resuspending cell pellets in 1 ml of 80% [v/v] ethanol supplemented with 1 µg/ml L-carnitine hydrochloride as internal standard and heating for 2 h at 60°C. After centrifugation at 17,000 x *g* for 5 minutes, the supernatant was transferred to a fresh vial and the pellet was again resuspended in 1 ml of 80% [v/v] ethanol and heated at 60°C for 2h. After centrifugation at 17,000 x *g* for 5 minutes, supernatants were combined and dried in a centrifugal evaporator.

Next, the dried extracts were dissolved in 1000 µL LC-MS grade water and filtrated through 0.2 µm filters (Omnifix-F, Braun, Germany). 1 µl of the cleared supernatants were analyzed using the high-performance liquid chromatograph mass spectrometer LCMS-8050 system (Shimadzu) and the incorporated LC-MS/MS method package for primary metabolites (version 2, Shimadzu) as described in (78).

### Bio-layer interferometry (BLI)

Protein interaction studies via Bio-layer interferometry (BLI) were performed on an Octet K2 instrument (Fortébio Molecular Devices (UK) Limited, Wokingham, United Kingdom), which allows simultaneous binding experiments on two channels, one of which is used as negative control. All experiments were done in buffer containing 50 mM Tris-HCl pH 7.4, 150 mM KCl, 2 mM MgCl_2_, 0.02% LDAO and 0.2mg/ml BSA. Effector molecules were used at following concentrations: 2 mM ATP, 2 mM 2-OG and 0.1, 0.25, 0.5, 1 and 2 mM for ADP. Interaction experiments were performed as reported previously (43). Briefly, His_8_-tagged variants of P_II_, namely P_II_(WT), P_II_(S49E), P_II_(I86N) and P_II_(ΔT)-His_8_ were used as ligands bound to Ni-NTA coated sensor tips. Various concentrations of GST-PirA from 125 to 12000 nM were used as analyte to display association reactions at 30°C. As preliminary experiments showed GST-PirA binds unspecifically to the Ni-NTA sensor tips, the non-interacting P_II_(ΔT) variant was used to saturate the tips and thereby remove unspecific binding. The binding of P_II_ ligands was performed by first loading 10 µg/ml of P_II_ on the Ni-NTA sensor tip, followed by dipping the tip into buffer for 60 s (to remove unbound P_II_) and recording the first baseline. To block unoccupied sites on the sensor surface, that cause disturbing unspecific binding, 72 µg/ml P_II_(ΔT) was loaded onto the tip. Afterwards, a second baseline was recorded for 60 s. Association and dissociation of analyte was carried out by dipping the tip first into GST-PirA solution for 180s and then transferring it into buffer solution for further 120s. In every single experiment, one sensor loaded with PII(ΔT) was used as negative control. To investigate the binding of GST tag alone and PirA without the GST-tag to P_II_ protein, parallel experiments were performed in the presence of 2 mM ADP. The interaction curves were achieved by subtracting the control curve and adjusting them to the average of baseline and dissociation steps. In every set of experiment K_D_ values were calculated by plotting concentration versus maximum response.

His_8_-tagged variants of P_II_ were prepared as previously described (79). For the preparation of recombinant PirA protein for BLI analysis, the *pirA* gene was cloned into XhoI and EcoRI sites of pGEX-4T-3 vector (GE Healthcare Life Sciences, Freiburg, Germany), encoding recombinant PirA with N-terminal-fused GST-tag. In addition, *pirA* was cloned into SapI-site of pBXC3GH vector (Addgene, Toddington,U.K.), encoding PirA with a C-terminally fused GFP-His_10_-tag. The plasmids were overexpressed in *E. coli* strain BL21(DE3). Purification of PirA with GST or GFP-His_10_ tags was performed as previously described (80, 81). To remove the GFP-His_10_-tag from PirA, 2.4 mg of recombinant PirA in 750 µl of PirA-buffer (50mM Tris/HCl, 100mM KCl, 100mM NaCl, 5mM MgCl_2_, 0.5mM EDTA, 1mM DTT, pH 7.8) were treated with 50 µl 3C protease (0.1. mg) at 4°C over night. Afterwards, 200 µl Ni-NTA agarose beads (QIAGEN GmbH, Hilden, Germany) were added to the mixture and gently shaken at room temperature for 60 min. The beads were removed by filtration and the tag-free PirA protein was dialyzed against PirA buffer containing 50% (v/v) glycerol and stored at −20° C until use.

## Acknowledgments

The project was funded by grants from the German Research Foundation (DFG) to SK (KL 3114/2-1), KF (Fo195/17-1) and MH (HA 2002/23-1), grants BIO2016-75634-P and PID2019-104513GB-100 from Agencia Estatal de Investigación (AEI) to FJF and MIMP, and BIO-0284 Group from Junta de Andalucía, all co-financed by FEDER (European regional development fund). The LC-MS/MS equipment at University of Rostock was financed through the HBFG program (GZ: INST 264/125-1 FUGG to M.H.). We also acknowledge the use of the facilities of the Centre for Biocatalysis (MiKat) at the Helmholtz Centre for Environmental Research (UFZ). The UFZ is supported by the European Regional Development Funds (EFRE, Europe funds Saxony) and the Helmholtz Association.

### Appendix

**Supplementary Table S1:**
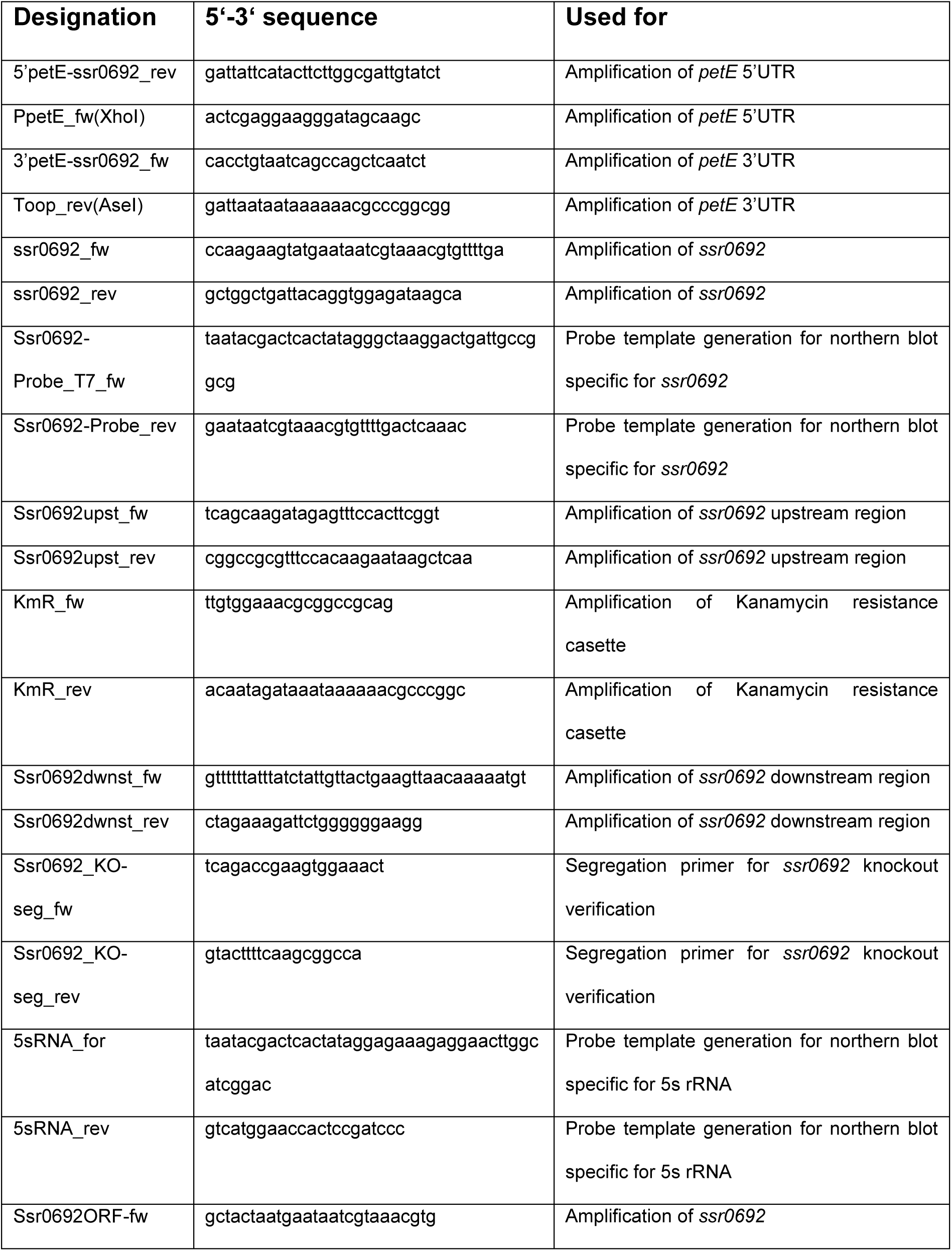
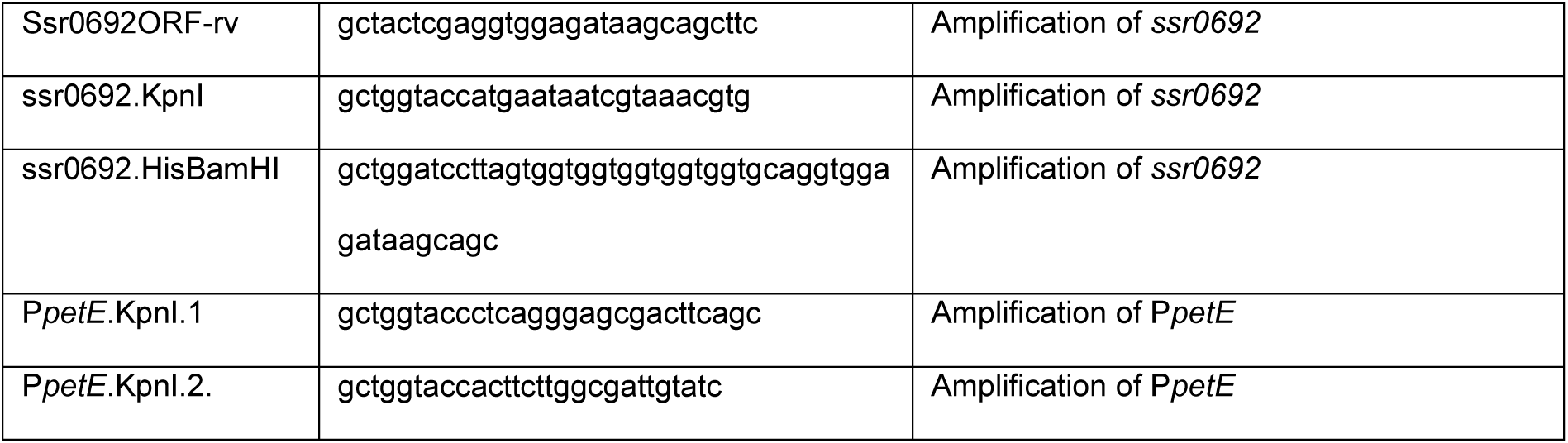
Primers used in this study.

**Supplementary Table S2:** Strains used in this study.

| Strain name | Source | Parental strain | Features |
| --- | --- | --- | --- |
| WT | Norio<br>Murata<br>(Jap.) | - | *Synechocystis* sp. PCC 6803, wild type, glucose-tolerant, non-motile |
| Δ0692 | This study | *Synechocystis* sp. PCC 6803 wild type | recombinant *Synechocystis* strain in which the *pirA* gene was deleted and replaced by a kanamycin resistance cassette |
| 0692<sup>+</sup> | This study | *Synechocystis* sp. PCC 6803 wild type | recombinant *Synechocystis* strain carrying the pVZ322-*PpetE::ssr0692* plasmid used for ectopic, Cu<sup>2+</sup>-inducible *pirA* expression |
| Δ0692 + *PpetE::ssr0692* | This study | Δ0692 | knockout strain for *pirA* with a *PpetE::ssr0692* construct inserted into the chromosome |

**Supplementary Figure S1:**
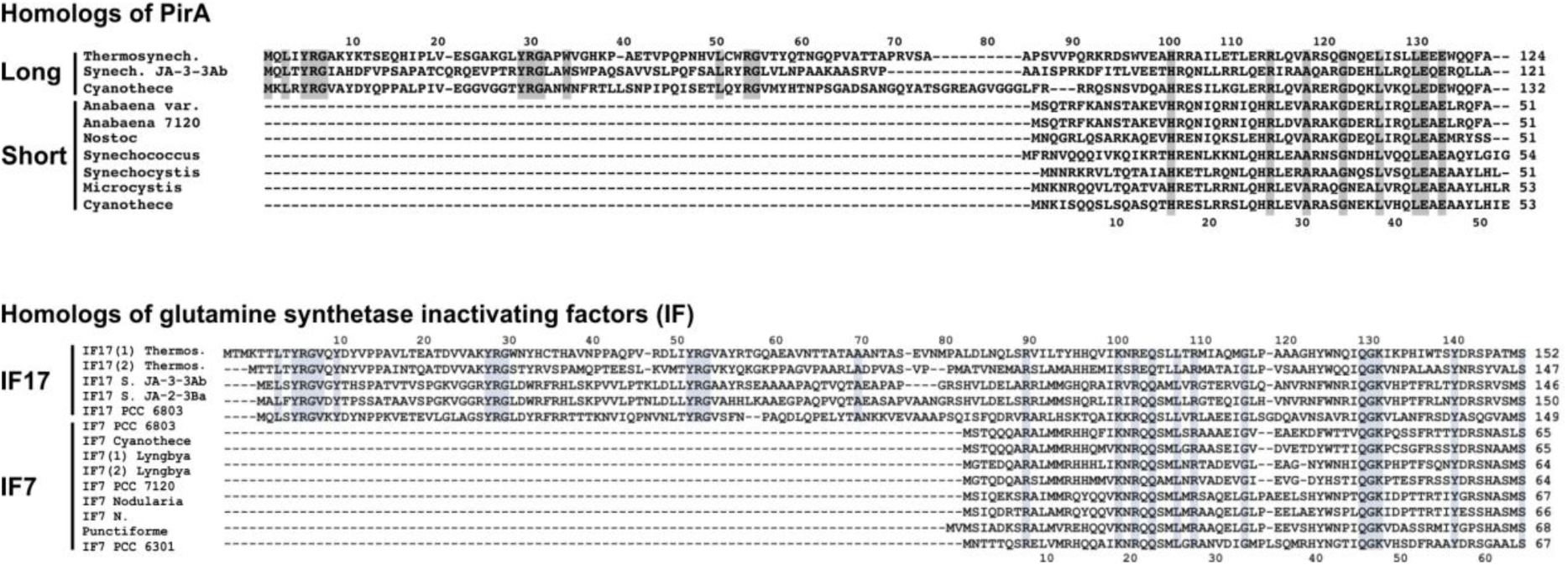
Alternative PirA variants. BLASTP analyses suggest alternative PirA versions with an extended N-terminal end in few strains. This resembles the glutamine synthetase (GS) inactivating factors (IF) all of which show a homologous C-terminus but only some show an N-terminal extension. These extensions show conserved amino acid residues that confer stability to the proteins that carry it (18). Those residues are also present in the N-terminal part of alternative PirA versions.

